# Intra-familial phenotypic heterogeneity and telomere abnormality in von Hippel-Lindau disease

**DOI:** 10.1101/526913

**Authors:** Jiangyi Wang, Xiang Peng, Cen Chen, Xianghui Ning, Shuanghe Peng, Teng Li, Shengjie Liu, Baoan Hong, Jingcheng Zhou, Kaifang Ma, Lin Cai, Kan Gong

## Abstract

Von Hippel-Lindau (VHL) disease is a hereditary cancer syndrome with poor survival. The current recommendations have proposed uniform surveillance strategies for all patients, neglecting the obvious phenotypic varieties. In this study, we aim to confirm the phenotypic heterogeneity in VHL disease and the underlying mechanism. A total of 151 parent-child pairs were enrolled for genetic anticipation analysis, and 77 sibling pairs for birth order effect analysis. Four statistical methods were used to compare the onset age of patients among different generations and different birth orders. The results showed that the average onset age was 18.9 years earlier in children than in their parents, which was statistically significant in all of the four statistical methods. Furthermore, the first-born siblings were affected 8.3 years later than the other ones among the maternal patients. Telomere shortening was confirmed to be associated with genetic anticipation in VHL families, while it failed to explain the birth order effect. Moreover, no significant difference was observed for overall survival between parents and children (p=0.834) and between first-born patients and the other siblings (p=0.390). This study provides definitive evidence and possible mechanisms of intra-familial phenotypic heterogeneity in VHL families, which is helpful to the update of surveillance guidelines.

## Introduction

Von Hippel-Lindau (VHL) disease (MIM 193300) is a rare autosomal dominant cancer syndrome caused by germline mutations in the VHL tumor suppressor gene (Latif *et al*. 1993; Lonser *et al*. 2003). Generally, the first VHL-related manifestation occurred in the second decade of patient’s life, and the penetrance is more than 90% by 70 years old (Ong *et al*. 2007; Nordstrom-O’Brien *et al*. 2010). Patients may develop various tumor types from early childhood through adulthood, including central nervous system hemangioblastomas (CHB), retinal hemangioblastomas (RA), clear cell renal cell carcinoma (RCC), pancreatic cyst and neuroendocrine tumors (PCT), pheochromocytomas (PHEO), endolymphatic sac tumors (ELST), epididymal and broad ligament cystadenomas (Walther and Linehan 1996; Lonser *et al*. 2003; Lonser *et al*. 2004; Butman *et al*. 2008).

The *VHL* gene is located on chromosome 3p25–26 and codes for VHL protein (pVHL), which is one of the most important tumor suppressor proteins (Gossage *et al*. 2015). pVHL forms an E3 ligase complex with elongation factor C and B (elongin C/B), cullin2 (CUL2) and RING finger protein (RBX1), and is critical for the ubiquitination and proteasomal degradation of HIF-αand other proteins (Gossage *et al*. 2015). The accumulation of HIF-α caused by mutations of *VHL* gene and dysregulation of several downstream proangiogenic factors have been identified to be the main cause for tumorigenesis in VHL disease. In recent studies, pVHL has also been found to act as a component of the ataxia telangiectasia mutated (ATM) dependent DNA-damage response (DDR) network, inactivation of which contributes to the genomic instability associated with sporadic RCC (Metcalf *et al*. 2014). Furthermore, ATM was reported to play a dual role in the sophisticated surveillance mechanisms at DNA double-strand breaks (DSBs) and in telomere regulation (Di Domenico *et al*. 2014). Thus, some HIF independent and DDR related pathways may play an important role in the pathogenesis of VHL related tumors.

Patients with VHL, a hereditary cancer syndrome, often ended with poor survival due to the complex phenotype and unclear pathogenesis (Wilding *et al*. 2012). Clinically, the mean life expectancy of VHL patients was reported to be 49 years old in a large cohort 30 years ago. Although there is no cure for the hereditary cancer syndrome, early diagnosis has made the median survival age improve to over 60 years old based on the establishment of active surveillance plans (Wang *et al*. 2018). However, the same plan was carried out for all the VHL patients according to the current surveillance recommendations, regardless of the fact that VHL disease demonstrates obvious phenotypic heterogeneity, which resulted in diagnostic and therapeutic delay of patients with early onset age. To provide personalized surveillance strategy for different individuals, it is a challenging issue to explore the phenotypic diversity in VHL disease. For patients in different families, genotype-phenotype correlations have been well constructed for years: patients with missense mutations are more likely to be affected by pheochromocytoma, while patients with truncating mutations confer a higher risk for RCC and CHB (Gallou *et al*. 2004; Ong *et al*. 2007). Nevertheless, the existing genotype-phenotype correlations do not work in VHL patients within the same family. Patients with consanguinity may develop tumors at different age even in the same family with the same genotype, implying that intra-familial phenotypic heterogeneity may be an important part in the complexity of VHL disease.

Genetic anticipation (GA) is one of the most common intra-familial phenotypic varieties for hereditary diseases, in which case the next generations are affected at an earlier age or manifest more serious presentations than their parents. Recently, GA has been reported in hereditary cancer syndromes including Lynch syndrome, Li-Fraumeni syndrome, dyskeratosis congenital and hereditary breast cancer, and was associated with genomic instability caused by shortening of telomere length in the next generations (Vulliamy *et al*. 2004; Tabori *et al*. 2007; Martinez-Delgado *et al*. 2011; von Salome *et al*. 2017). However, studies on GA for VHL disease were rare. In our previous study, we for the first time found that telomere shortening was associated with GA in Chinese patients with 18 VHL families (Ning *et al*. 2014). Nevertheless, Laura reported that telomere abnormalities might just be primarily somatic in origin rather than a cause of GA in 20 families, making the conclusion ambiguous (Aronoff *et al*. 2018). Considering the small number of families enrolled in the two studies, only the anticipation of the first tumor was reported rather than separate analysis about the same tumor between different generations. Thus, further study on intra-familial phenotypic heterogeneity and the underlying mechanism is needed for better understanding of the disease.

In our present study, we recruited 80 unrelated VHL families and analyzed the phenotype between successive generations and among siblings in the same generation. The results showed that the average onset age was 18.9 years earlier in children than that in their parents, and the first-born siblings were affected by VHL-associated tumors 5.6 years later than the other ones, implying the existence of genetic anticipation and birth order effect within VHL families. Furthermore, we found that telomere shortening in the next generation was associated with genetic anticipation in VHL families, giving a clue that genomic instability might play an important role in the pathogenesis of VHL-associated tumors. The results are helpful for genetic counseling and future updates from the present uniform recommendations to individualized surveillance plan.

## Materials and methods

### Editorial Policies and Ethical Considerations

This study was approved by the Medical Ethics Committee of Peking University First Hospital (Beijing, China) and written informed consent was obtained from all subjects.

### Patients and samples

From 2009 to 2016, a total of 348 patients from 133 families were diagnosed with VHL disease at the Institute of Urology, Peking University according to the clinical criteria and VHL gene detection as previously described (Wang *et al*. 2017; Liu *et al*. 2018). All the patients diagnosed by clinical symptoms had at least one affected relative identified by VHL mutation test, so that their genotype could be predicted. Forty-eight patients were excluded because of uncertain clinical information, and another 24 patients were excluded because they did not have affected parents, children or siblings. Therefore, a total of 276 patients from 80 families were enrolled in this study. Patients in the different generations formed 151 parent-child pairs for genetic anticipation analysis, and patients with the identical parent formed 77 sibling pairs for birth order effect analysis. Onset age was defined as the age when the first symptom or sign occurred. The basic clinical data was shown in Table S1. Among all the individuals enrolled, 141 patients with available peripheral blood leukocyte DNA sample were brought into analysis for the relationship between relative telomere length and phenotypic diversity.

### Relative telomere length measurement

Genomic DNA was extracted from peripheral blood leukocyte using a blood DNA extraction kit (Tiangen Biotech). Quantitative real-time PCR was used to measure telomere length as described by Cawthon (Cawthon 2002). The PCR reaction was run in7500 instrument (Applied Biosystems), containing 5 μL 2X SYBR master mix (Takara), 30ng genomic DNA, 300nmol/L telomere primer Tel1 and 900 nmol/L Tel2, or 200 nmol/L single copy gene primer 36B4u and 500 nmol/L 36B4d. The primer sequences and PCR reaction procedure were described in our previous study (Wang *et al*. 2017). Whenever possible, samples from different groups were run on the same plate. A standard curve was constructed to assess the amplification efficiency (E) using a control DNA sample (male, 45 years old) diluted by 1/4 serial from 50ng to 0.19ng, and the same sample was detected in every batch of PCRs as the inter-run calibration. The telomere repeats (T) was described by (E_Tel,Sample_)^−Ct(Tel,sample)^/(E_Tel,Calibrator_)^−Ct(Tel,calibrator)^, and the copy number of 36B4 (S) was (E_36B4,sample_)^−Ct(36B4,sample)^/(E_36B4,calibrator_)^−Ct(36B4,calibrator)^. Relative telomere length (RTL) was calculated by T/S. Age-adjusted RTL (aRTL) was obtained based on the telomere-age curve constructed in our study before (Ning *et al*. 2014).

### Statistical Analysis

Patients’ entire lifetimes from birth until death or the end of follow-up in Dec 2016 were included for analysis, and the median observation time was 39.3 years/person (range 1–75 years) with a total 10860 person-years. The survival of patients and age-related penetrance of the five VHL-associated tumors were analyzed using Kaplan-Meier plots and log-rank test. For the analysis of genetic anticipation and birth order effect, we first used paired t-test to examine the difference of onset age between affected patient pairs. Considering the truncating bias caused by clinical data in paired t-test, we confirmed the results using the nonparametric method of Rabinowitz and Yang 1 (RY1) and nonparametric method of Rabinowitz and Yang 2 (RY2) which were special statistical methods for genetic anticipation analysis (Boonstra *et al*. 2010). Furthermore, the Cox proportional hazards model (CPH) was used to evaluate the effect of generation on phenotype when we included both affected and unaffected patients (Boonstra *et al*. 2010).

Statistical analyses were performed using SPSS13.0 and R software, and a two-sided p-value of less than 0.05 was considered to be statistically significant.

### Data Availability

The authors affirm that all data necessary for confirming the conclusions of the article are present within the article, figures, tables and supplementary files.

## Results

### Genetic anticipation in different generations

Genetic anticipation was analyzed in 151 parent-child pairs from 80 VHL families. As we reported before, the age-related penetrance in children was obviously higher than that in their parents (p<0.001) (Figure1), and the result was not affected by mutation origin and mutation type (Figure S1). In this study, we further analyzed the five common VHL-related tumors CHB, RA, RCC, PCT and PHEO, respectively. The results showed that children had higher age-related penetrance in all five tumors than their parents (Figure1). To further support our results, we assessed the anticipation in 20 grandparents-parents-children groups from 16 families. The data revealed that patients showed higher age-related penetrance in successive generations (p<0.001) (Figure 1).

**Figure 1.**
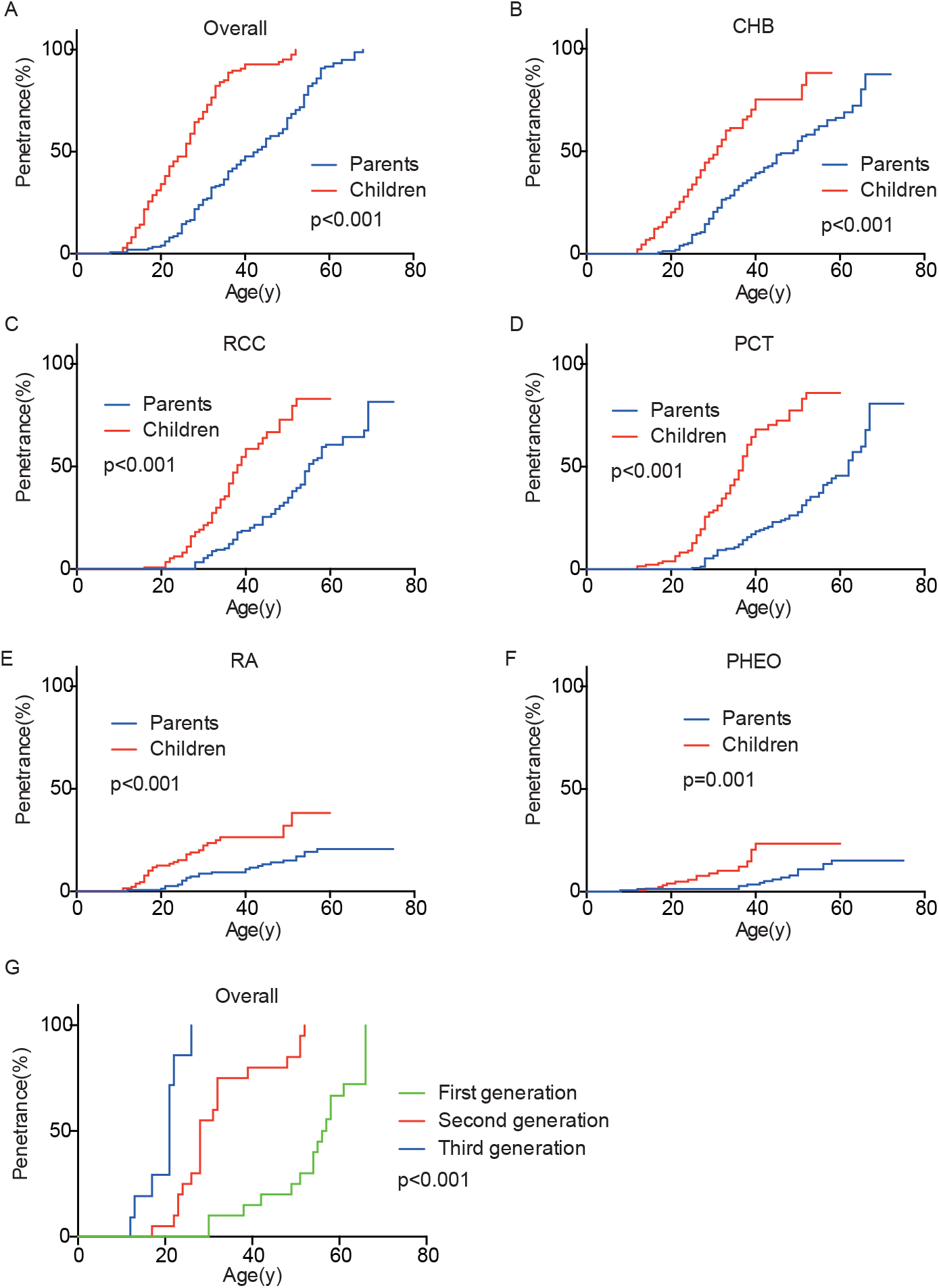
Genetic anticipation between parents and children in different generations. Children showed higher age-related risk than their parents for overall tumors (A) and CHB (B), RCC (C), PCT (D), RA (E), PHEO (F), respectively. Age-related penetrance was also assessed in 16 families with three generations and showed the same tendency (G).

To evaluate the exact difference of onset age between parents and children, we analyzed the 114 tumor-affected parent-child pairs and found that the average onset age was 18.9 years earlier in children (Table 1) (p<0.001), which was 2.1 years longer than we previously reported in 34 parent-child pairs (Ning *et al*. 2014). Considering the possible bias caused by truncating data, the result was reanalyzed using RY1 and RY2 with the affected parents-children pairs and at last by CPH model with affected and non-affected VHL patients. As expected, the difference of onset age between parents and children was statistically significant in all the three methods (Table 1 and Table S2). For the onset age of the five common VHL-related tumors respectively, a similar tendency was observed (Table S3). The mean of onset age difference (MOAD) ranged from 17.2 years (CHB) to 34.6 years (PHEO) in different tumors.

**Table 1.** Genetic anticipation in affected parents-children pairs

|  | Mean onset<br>age (y) | Pairs | MOAD | Statistics |  |  |
| --- | --- | --- | --- | --- | --- | --- |
|  |  |  |  | Paired<br>t-test | RY1 | RY2 |
| **Parents** | 43.7±12.4 | 114 | 18.9 | 17.293<sup>***</sup> | 1.789<sup>*</sup> | 2.599<sup>**</sup> |
| **Children** | 24.7±9.7 |  |  |  |  |  |

\*\*\*p <0.001, \*\* p<0.01, \* p<0.05
Abbreviations: MOAD, mean of onset age difference.

### Birth order effect in the same generation

Given that the next generation showed a much earlier onset age than the first generation, we further assessed the phenotypic heterogeneity in the same generation, which was called the birth order effect. A total of 77 sibling pairs from the identical parents were analyzed, and the distribution of onset age similarly displayed a “shift to left” phenomenon in younger siblings (p=0.036) (Figure 2). On average, the first-born siblings were affected by VHL-associated tumors 5.6 years later than the other ones, which was statistically significant in paired t-test (p<0.001), RY1 (p<0.05) and RY2 (p<0.05) (Table 2). Interestingly, when we analyzed the data only in the paternal sibling pairs, the birth order effect disappeared (Figure 2 and Table 2). In contrast, in the maternal sibling pairs, the MOAD increased to 8.3 years later between the first-born siblings and others (Figure 2 and Table 2).

**Figure 2.**
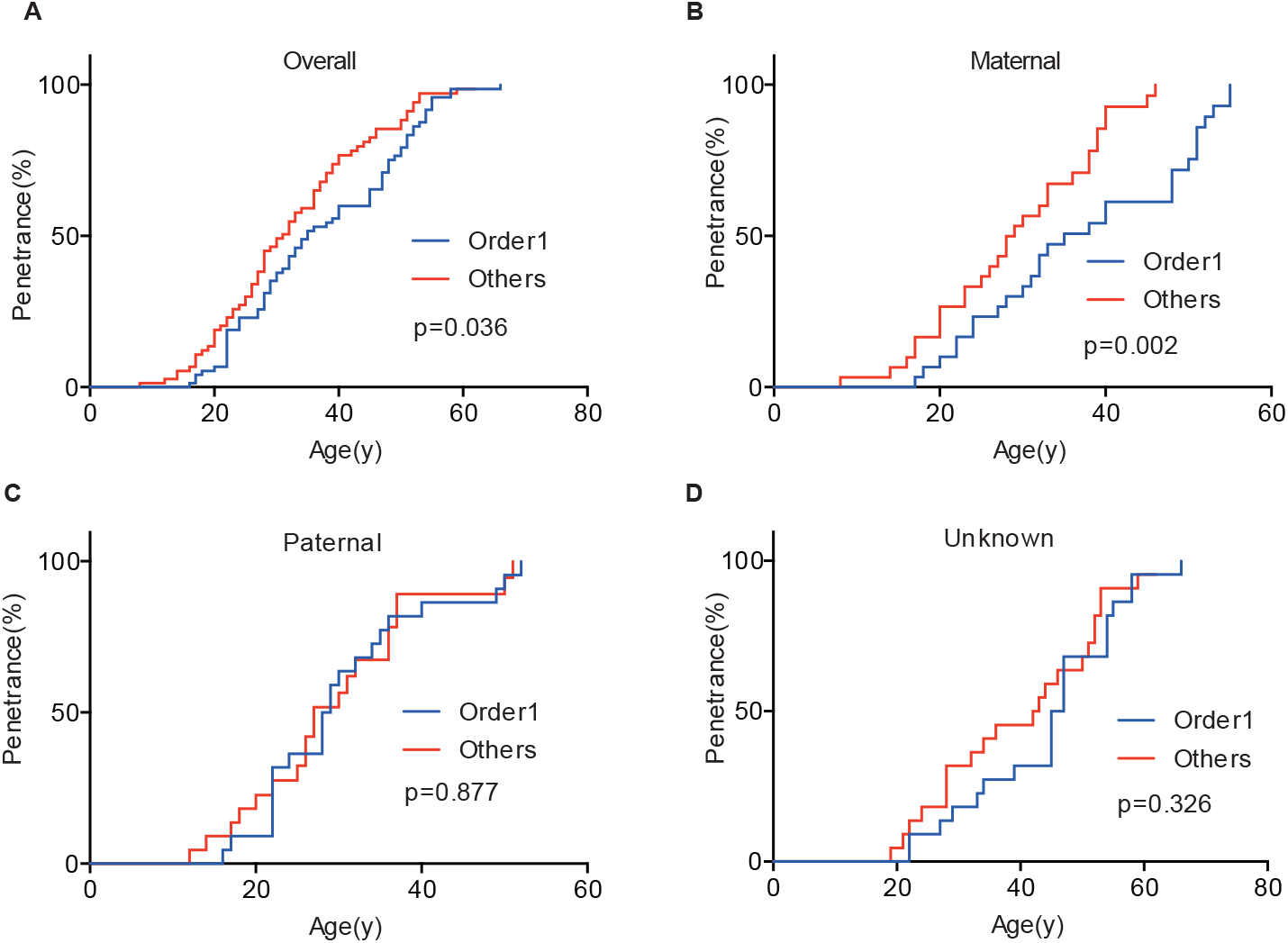
Birth order effect between the first-born siblings and the others in the same generation. The first-born siblings displayed lower age-related penetrance than the others for overall tumors (A). Among siblings with an identical affected mother, the birth order effect became more evident (B). When the mutated VHL allele was from fathers or the origin was not clear, the effect disappeared (D).

**Table 2.** Birth order effect in affected offspring

|  | Pairs | Mean onset age (y) |  | MOAD(y) | Statistics |  |  |
| --- | --- | --- | --- | --- | --- | --- | --- |
|  |  | Older | Younger |  | t-test | RY1 | RY2 |
| **Overall** | 69 | 37.5±12.5 | 31.9±11.7 | 5.6 | 3.316<sup>***</sup> | 1.897<sup>*</sup> | 2.303<sup>*</sup> |
| **Maternal** | 28 | 37.3±12.8 | 29.0±9.7 | 8.3 | 3.175<sup>**</sup> | 1.706<sup>*</sup> | 2.113<sup>*</sup> |
| **Paternal** | 20 | 31.4±10.0 | 28.7±10.6 | 2.7 | 1.326 | 0.707 | 1.693 |

\*\*\* p<0.001, \*\* p<0.01, \* p<0.05
Abbreviations: MOAD, mean of onset age difference

### Telomere length and intra-family phenotypic diversity in VHL disease

The relationship between telomere length and intra-family phenotypic diversity for VHL disease was next investigated. A total of 55 parent-child pairs with data of telomere length were enrolled for analysis, and the result showed that in 80% (44/55) pairs, children had significantly shorter aRTL than their parents (p<0.001) (Figure 3). Furthermore, in the 33 parent-child pairs with clear onset ages, 28 children had younger onset age with shorter aRTL, while their parents had older onset age with longer aRTL (Table 3). Among the 16 families with three generations, only 3 families were available for DNA samples of three generations (Family 42, 48, 66), and all of them displayed a tendency of shorter telomere length along with generations (Figure S2). With regard to the sibling pairs, 16 out of 25 pairs showed relatively shorter aRTL in the younger siblings, but the result was not statistically significant (p=0.585) (Figure 3). In the 19 tumor-affected sibling pairs, only 8 younger patients had shorter aRTL with younger onset age (Table S4). The results indicated that telomere shortening was associated with genetic anticipation in VHL families, while the association between telomere length and birth order effect might be uncertain.

**Figure 3.**
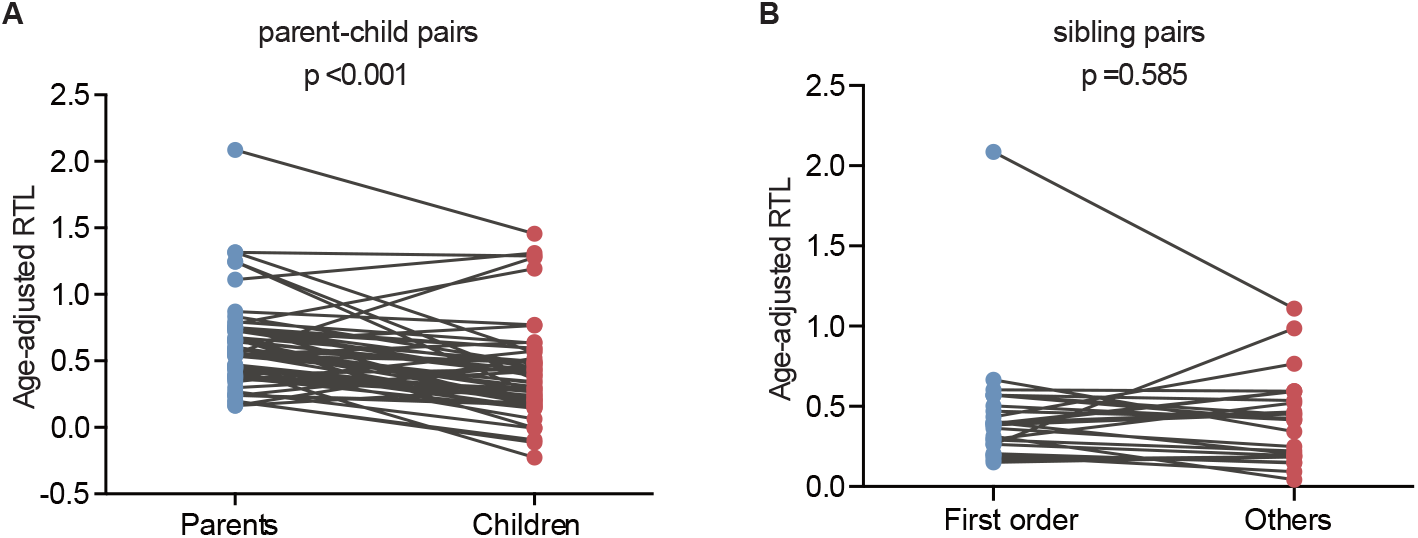
Age-adjusted relative telomere length in parents and children and in patients with different birth order. (A) Children showed a significant shorter telomere length than their parents. (B) No significant difference of telomere length was observed between the first-born sibling and the others.

**Table 3.** Telomere length between parents and children

|  | **Onset age (P>C)** | **Onset age (P≤C)** | **Overall** |
| --- | --- | --- | --- |
| **Age-adjusted RTL**<br>**(P>C)** | 28 | 2 | 30 |
| **Age-adjusted RTL**<br>**(P≤C)** | 2 | 1 | 3 |
| **Overall** | 30 | 3 | 33 |

Abbreviations: P, Parents; C, Children; RTL, relative telomere length

### Survival of VHL patients in different generations and birth order

In the 151 parent-child pairs, 58 parents and 10 children died. The median survival for parents was 62 years old, while the survival for children was undefined (Figure 4). No significant difference was observed for overall survival between parents and children (p=0.834) (Figure 4). Similarly, there was no difference between the overall survival for the first-born patients and the other siblings (p=0.390) (Figure 4). Considering the diagnostic and therapeutic improvement in the next generation and later-born siblings, the similar overall survival in some sense confirmed the genetic anticipation and birth order effect.

**Figure 4.**
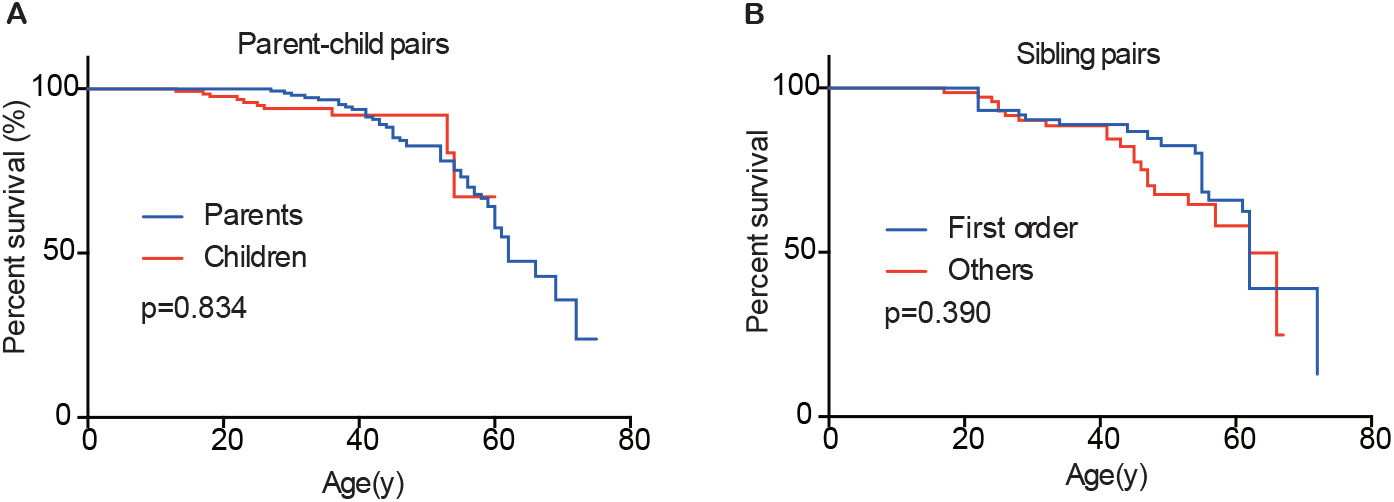
Overall survivals of patients in different generations and in different birth order. No significant difference was observed between parents and children (A) and between the first-born sibling and the others (B).

## Discussion

Studies on the average onset ages of VHL-related tumors have helped the VHL Alliance to propose the VHL tumor surveillance regimen for the last decades, according to which routine imaging screening should start at 1 year old for RA, 8 for PHEO, 16 for CHB and 8 for RCC and PCT.(Vhl 2015) Clinically, there exist several strategies of surveillance for VHL disease in the world, and the initial screening age differ from each other because of the different phenotypic features of patients they refer to (Hes *et al*. 2001; Binderup *et al*. 2013; Kruizinga *et al*. 2014; Rednam *et al*. 2017). Recently, our previous study and another Canadian study demonstrated that children in VHL families presented manifestations more than 10 years earlier than their parents, suggesting that surveillance plan for children should not only consider the regular VHL tumor surveillance regimen, but also the onset age of their parents (Ning *et al*. 2014; Aronoff *et al*. 2018). However, the limited number of patients involved in the studies weakened the reliability, and both of the two studies did not analyze the onset age of the five common VHL-related tumors CHB, RA, RCC, PCT and PHEO in different generations, respectively. In our present study, we confirmed the genetic anticipation phenomenon in VHL families, and interestingly, we also observed the birth order effect among VHL siblings from the same affected mother. This suggests that patients with affected parents or born in the later order within the family should start screening much earlier than others. Specifically, children in VHL families should receive MRI examination of brain and spinal cord from 12 years old rather than 14 to 16 years old recommended by the current strategies, since six children are affected by CHB before 14 (three for 12 years and three for 13) in our study. As to PHEO, we agree with the VHL Alliance and the regular abdominal imaging for children should start at 8 years old instead of 10 years recommended by Hes (Hes *et al*. 2001).

GA has been a controversial issue since it was first introduced because of the bias caused by data evaluation and statistical method the authors choose (Gruber and Mukherjee 2009). Paired t-test is the most common method used for anticipation analysis, but it may introduce a truncation bias, which will increase the type I error (Heiman *et al*. 1996). To lower the truncation bias caused by paired t-test, we further used two special statistical methods for genetic anticipation analysis (RY1 and RY2) in this study, reaching the same results. However, all the three methods are carried out with affected parent-child pairs, missing the data of unaffected patients. Thus, the CPH model was finally used to confirm GA with all the 151 parent-child pairs. As expected, the difference of onset age between parents and children is significant in all the four statistical methods.

GA and birth order effect have also been thought to be the myth caused by the improvement of diagnosis and surveillance in the younger patients, which is a source of ascertainment bias. People tend to take more active surveillance plan when they have relatives diagnosed with the hereditary disease, and this may lead to early detection of tumors before symptom. To reduce this kind of bias, we only analyzed the data of patient pairs presenting symptoms. It turned out that the differences of onset age between different generations and different birth orders were highly significant (Table 1, Table 2) (p <0.001). Moreover, children are affected by VHL related tumors before their parents present symptoms in 24% parent-child pairs, and similarly, about 38% siblings born in later order suffer tumors before the first-born patients become symptomatic. Thus, the ascertainment bias will not influence the final conclusion in our study.

To explore the molecular mechanism for genetic anticipation, we measured the age-adjusted relative telomere length of blood leukocytes in 10 parent-child pairs in our previous study and found that the age-adjusted telomere length was significantly shorter in a child than in his or her parent in all of the 10 parent-child pairs (Ning *et al*. 2014). Furthermore, VHL patients showed significantly shorter telomere length than healthy family controls, and a positive correlation was found between telomere length and onset age of the five major VHL related tumors, respectively (Wang *et al*. 2017). However, in another study from Canada on genetic anticipation with 15 VHL patients, only granulocyte telomeres from VHL patients were significantly shorter than those from healthy controls, and no significant difference was observed between different generations in the 6 parent-child pairs, implying that genetic anticipation in VHL is not caused by telomere abnormalities (Aronoff *et al*. 2018). The different results in the two studies above may be due to bias caused by the small sample size and different methods in measuring the telomere length. In this study, we evaluate the relationship between telomere shortening and genetic anticipation in a large VHL cohort and confirm that the age-adjusted relative telomere length was shorter in children than in their parents in 44 out of 55 parent-child pairs. As we all know, the telomeres in most cells shorten throughout human life, meaning that the telomere length would be influenced by age in which the blood sample was collected. Therefore, we use the age-adjusted relative telomere length (calculated as the difference between predicted normal relative telomere length at the DNA-obtained age and the relative telomere length actually measured) instead of relative telomere length in our analysis. The results confirm that telomere shortening is the molecular mechanism for genetic anticipation in VHL families.

Nevertheless, telomere shortening seems not to play a role in the mechanism of birth order effect in VHL disease. Although parous women have been reported to have shorter telomere length than nulliparous women (Pollack *et al*. 2018), we did not observe significant difference for telomere length between siblings in the first order and others. Currently, there exist three explanations for birth order effect, namely more infectious exposures in later-born children, a higher level of hormonal exposures in first pregnancies and a higher level of microchimerism in the later-born individuals(Von Behren *et al*. 2011). In this study, we find that birth order effect only exists in maternal sibling pairs, giving us a clue that the differences of hormonal exposures during pregnancies may contribute to the phenotypic diversity of VHL patients in different birth order. However, we don’t have enough data to confirm the hypothesis. The specific mechanisms for birth order effect in VHL families need further exploration.

In conclusion, this study provides definitive evidence of intra-familiar phenotypic variety in VHL families including genetic anticipation and birth order effect. Clinicians should take the position in the family tree into consideration when they are making surveillance plan for VHL patients. Although we confirm that telomere shortening plays a role in the molecular mechanism of phenotypic heterogeneity within VHL families, the detailed effect of inactivated VHL protein on genomic instability remains unclear, and the explanation for birth order effect needs further exploration.

## Acknowledgments

We thank Tingting Yan, Division of Gastroenterology and Hepatology, Renji Hospital, Shanghai, China, for her revision of the manuscript.

This work was supported by the National Natural Science Foundation of China (grant number: 81572506), the Fundamental Research Funds for the Central Universities (grant number BMU2018JI002), Beijing Municipal Science and Technology Commission (grant number Z151100003915126) and Beijing Municipal Commission of Health and Family Planning (grant number 2016-2-4074).

---

**Figure S1.**
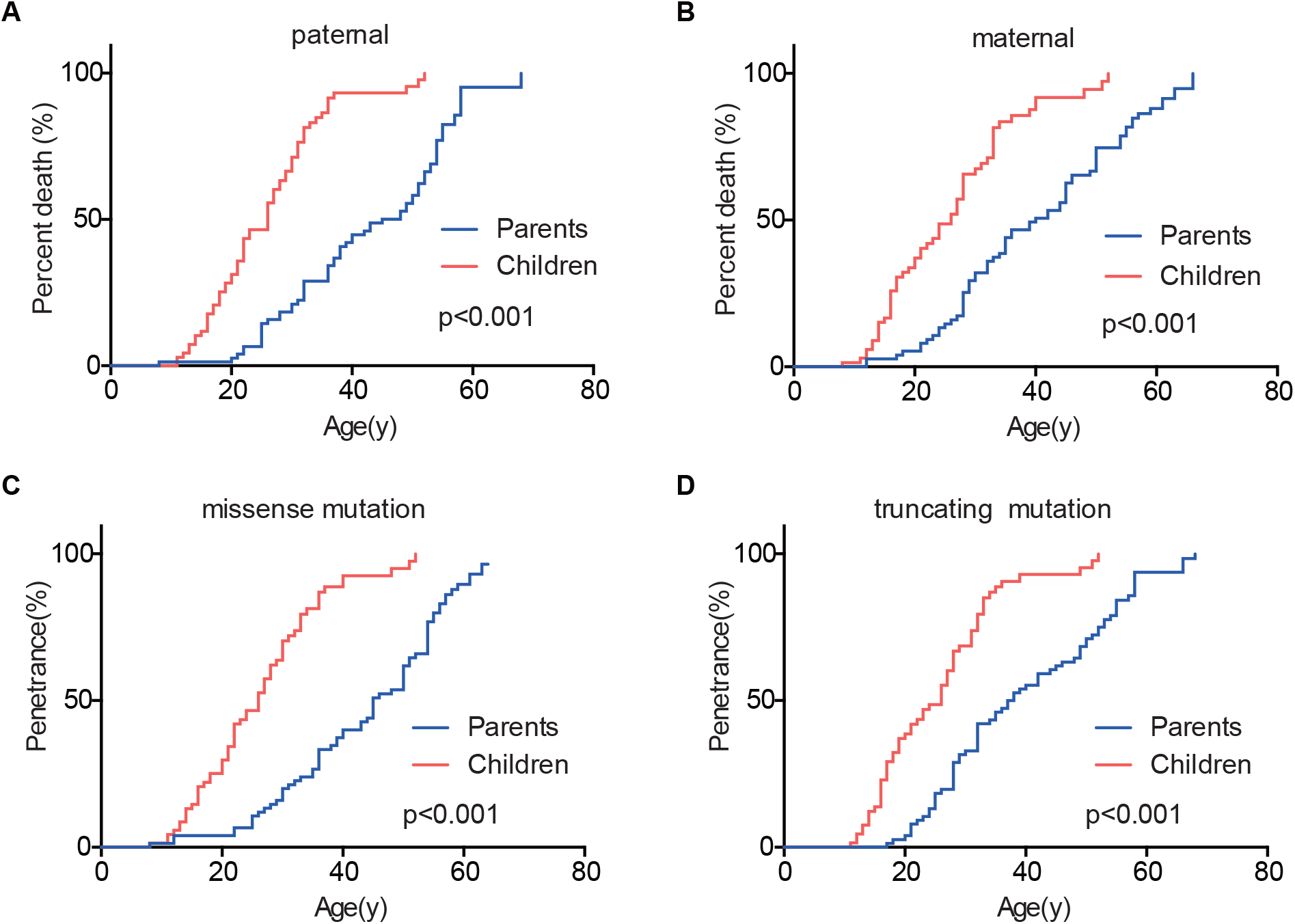
The effect of mutation origin and mutation type on genetic anticipation.

**Figure S2.**
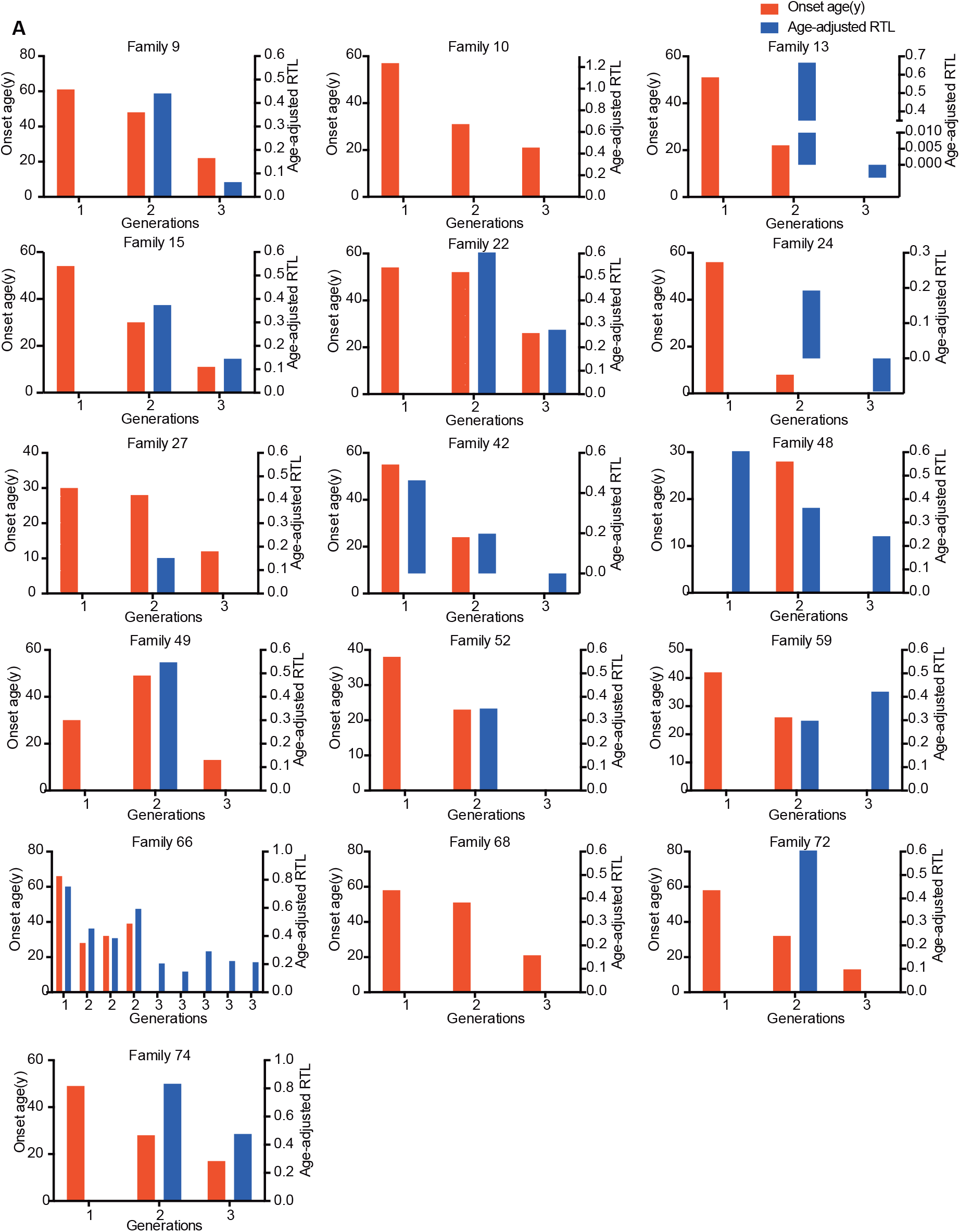
The relationship between genetic anticipation and telomere shortening in 16 VHL families with three generations.

**Table S1.**
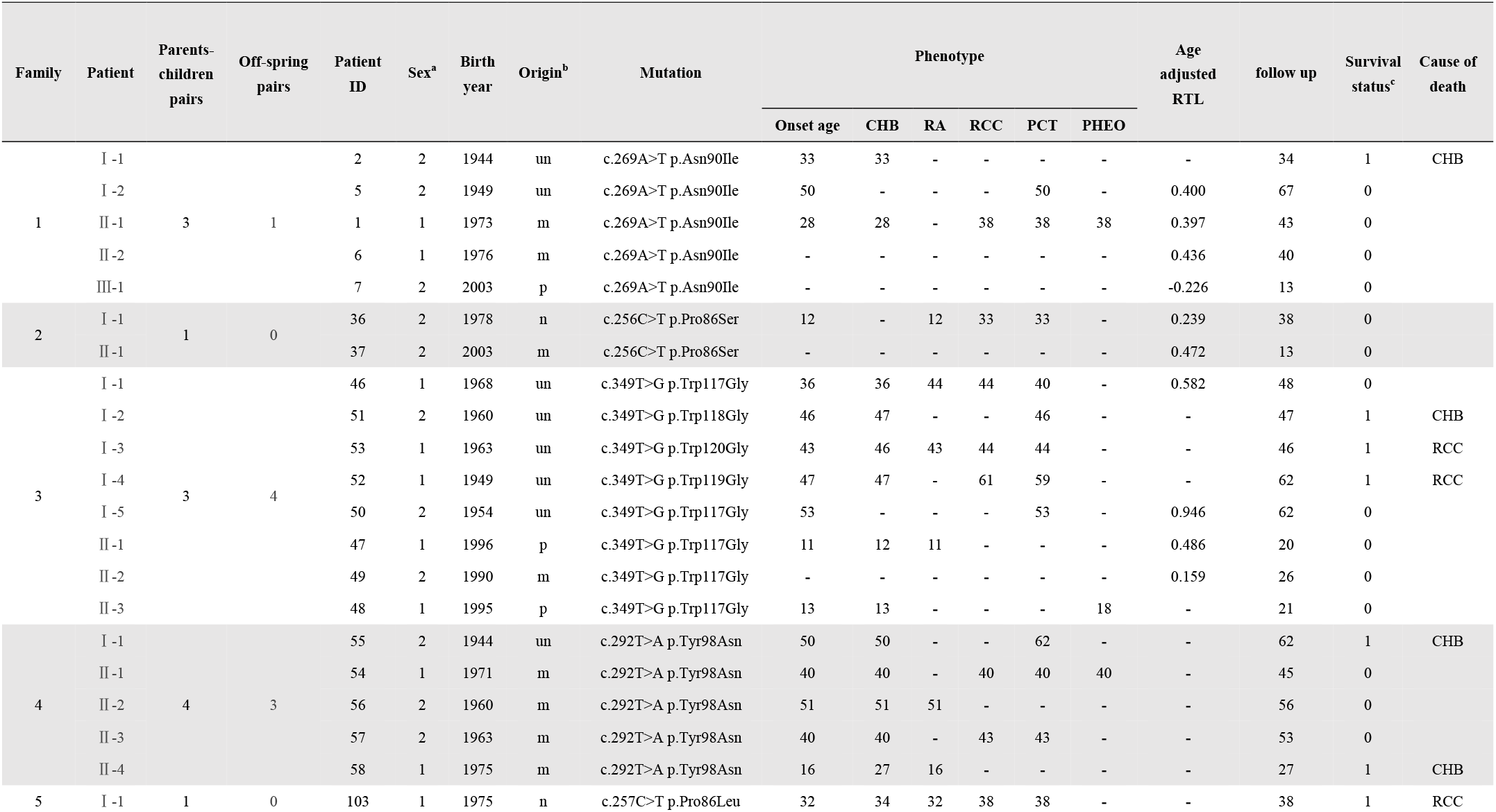
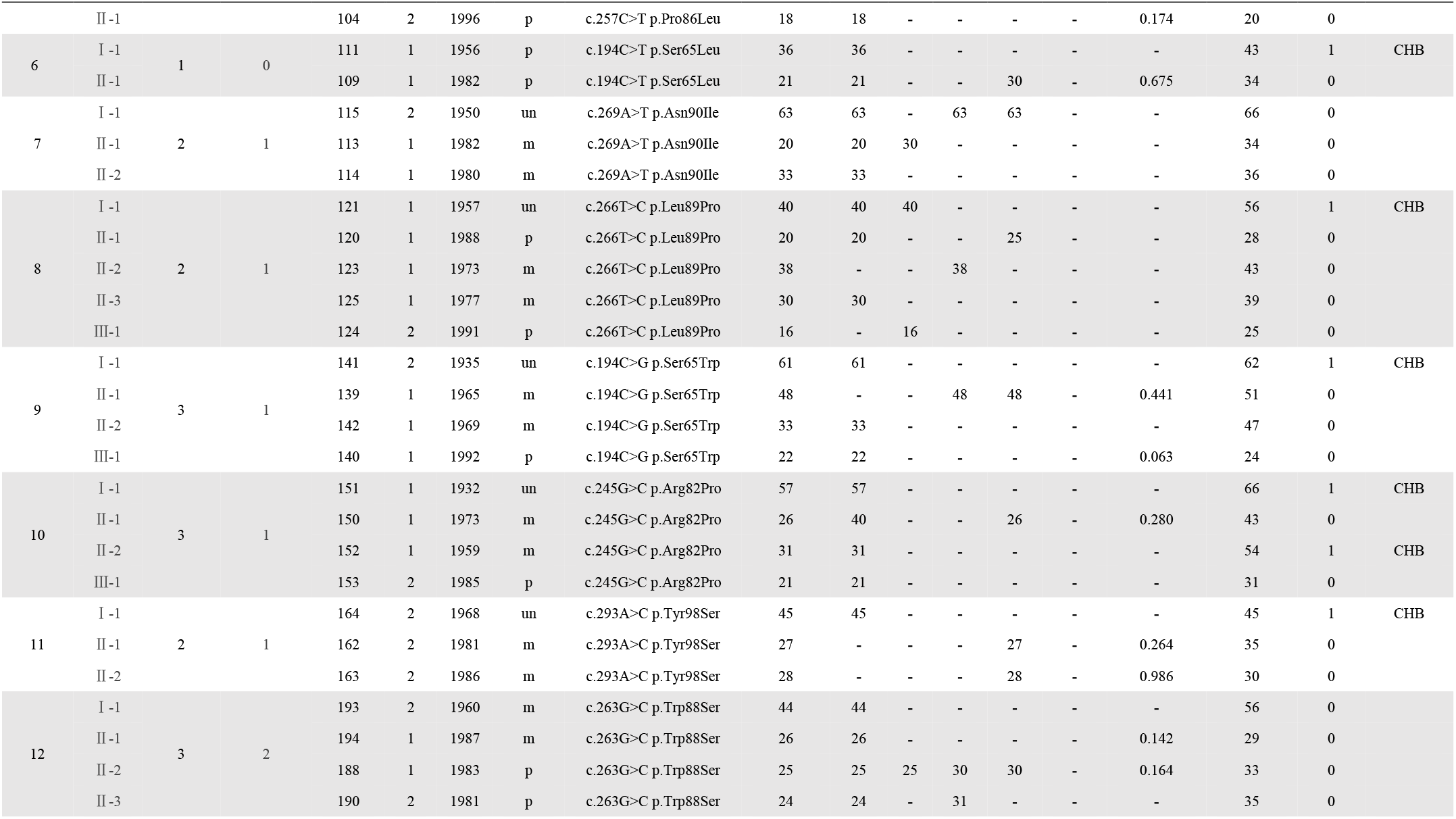
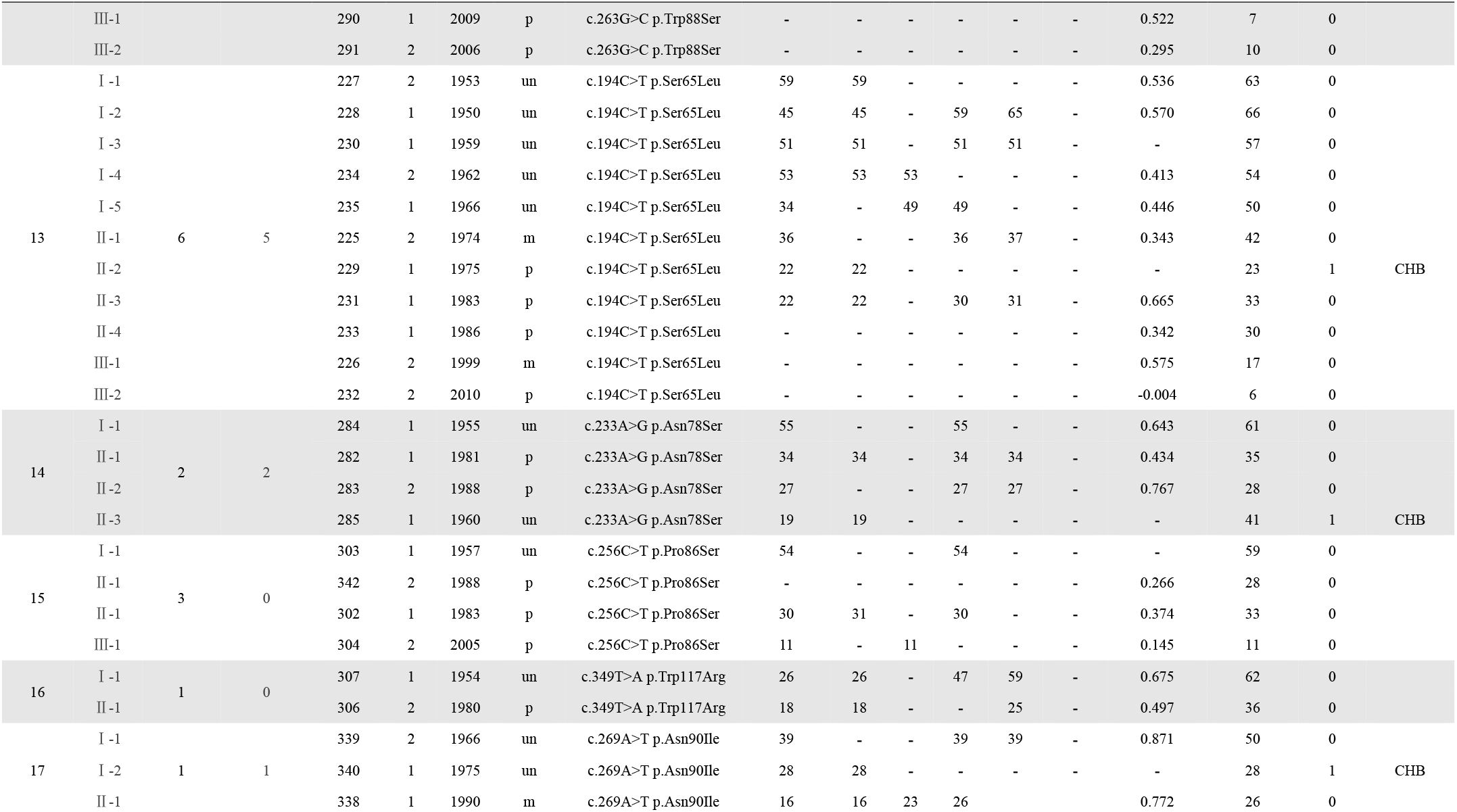
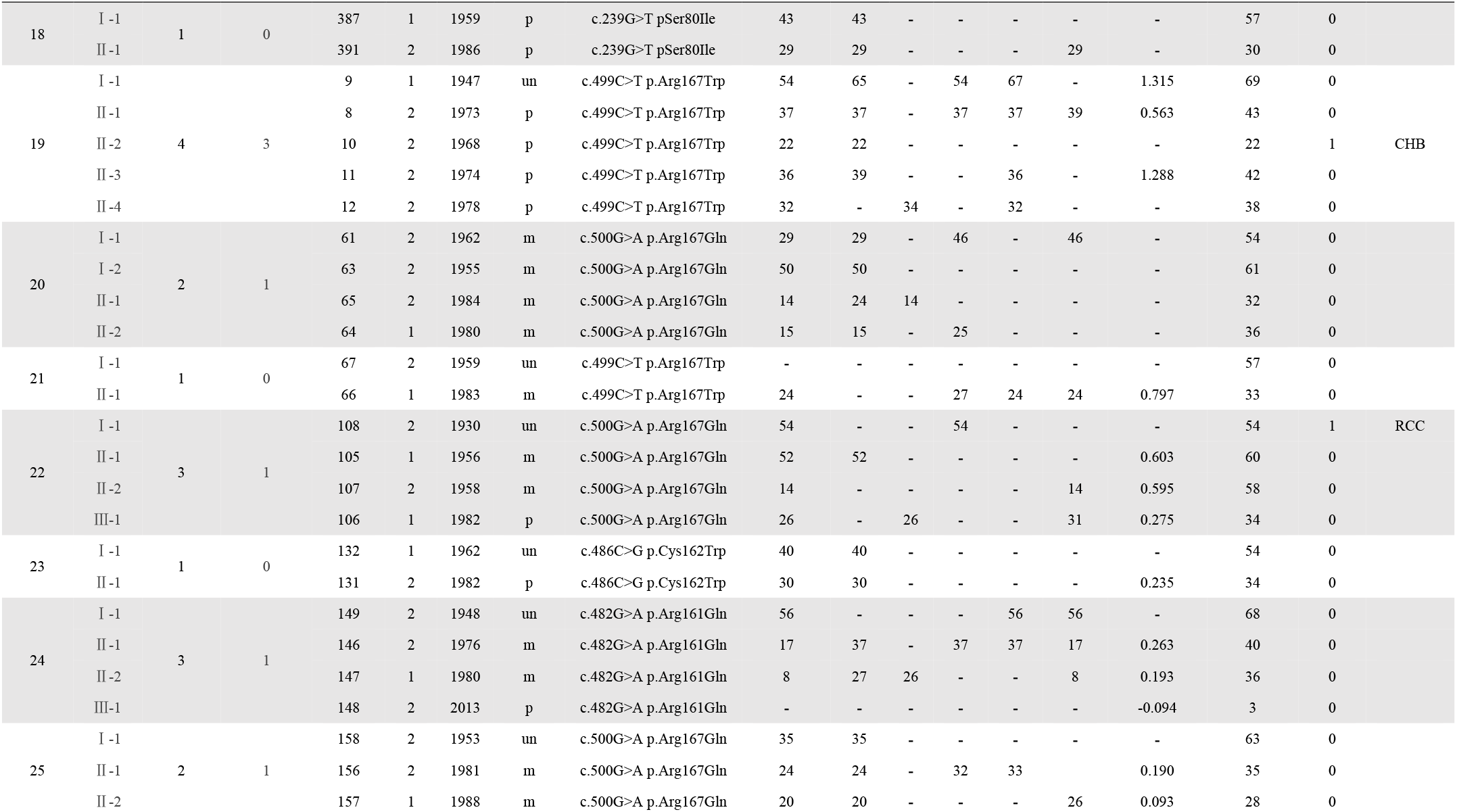
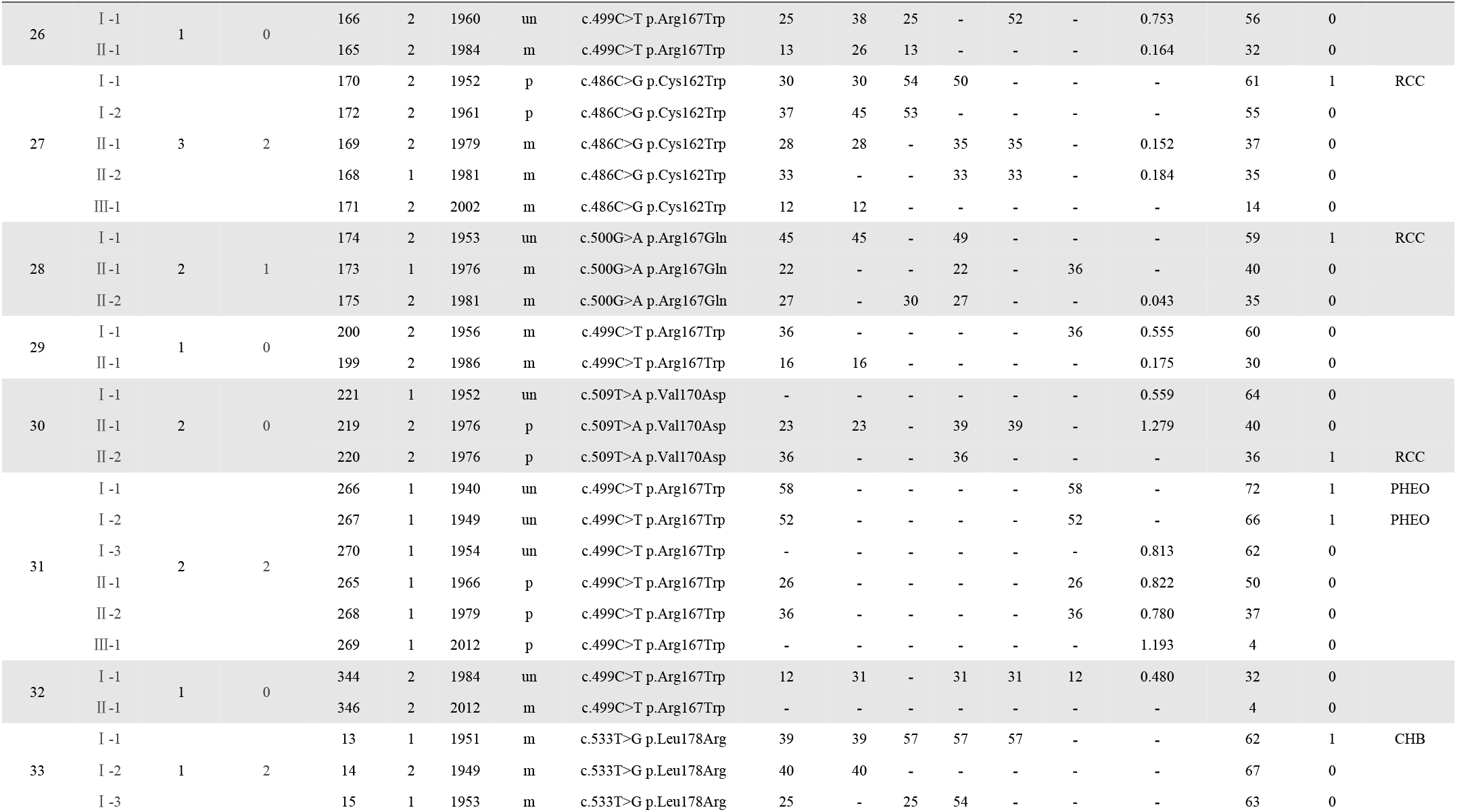
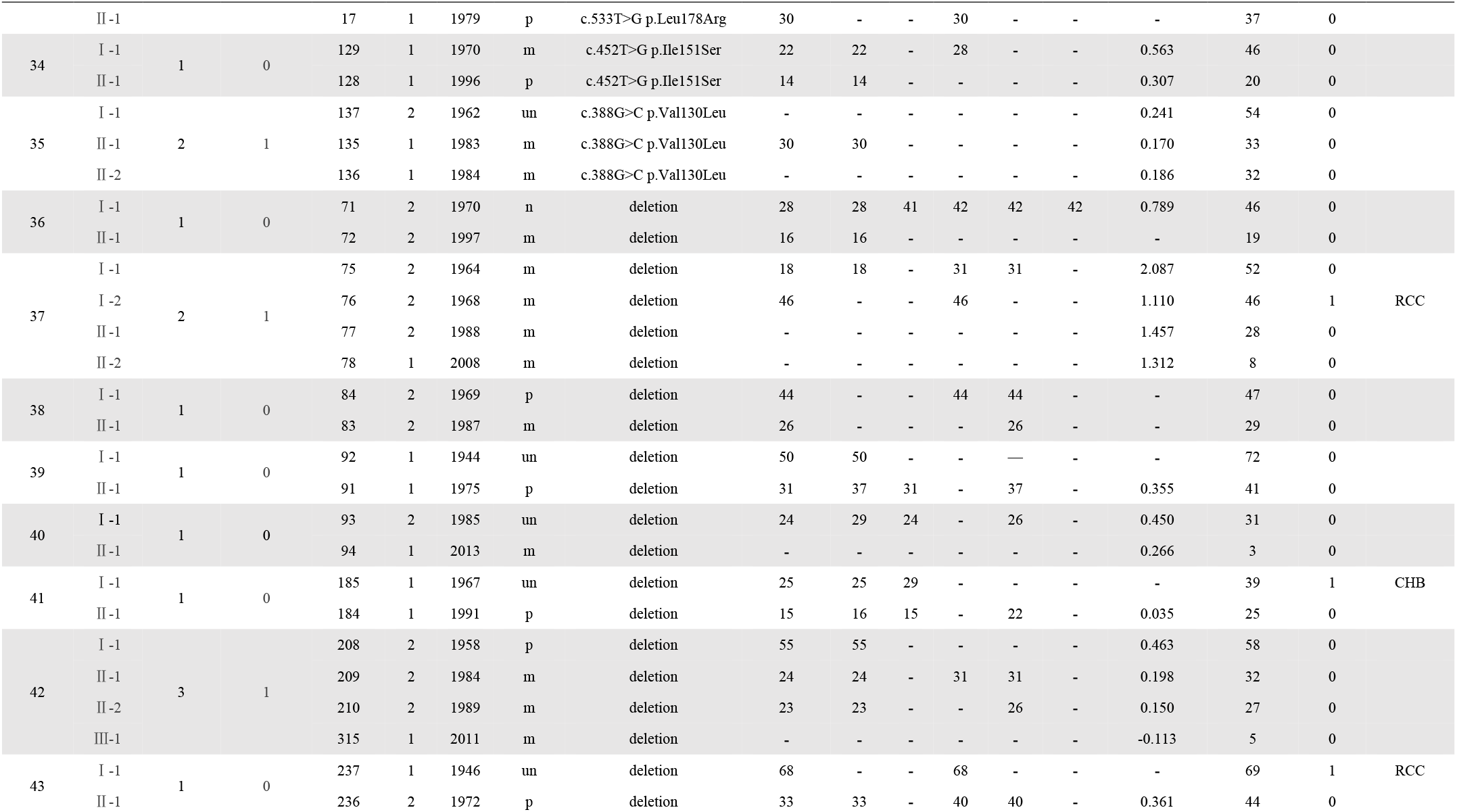
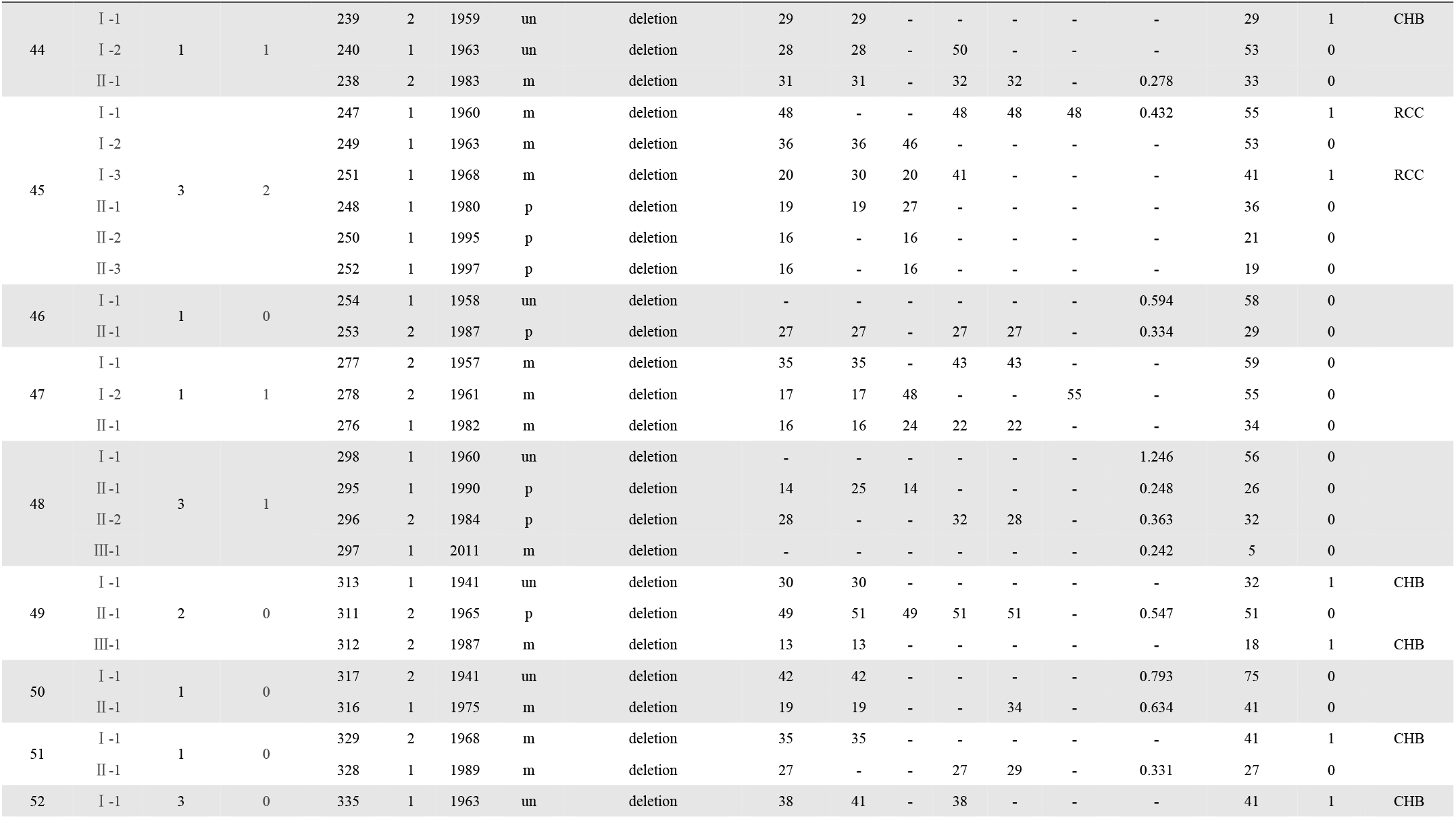
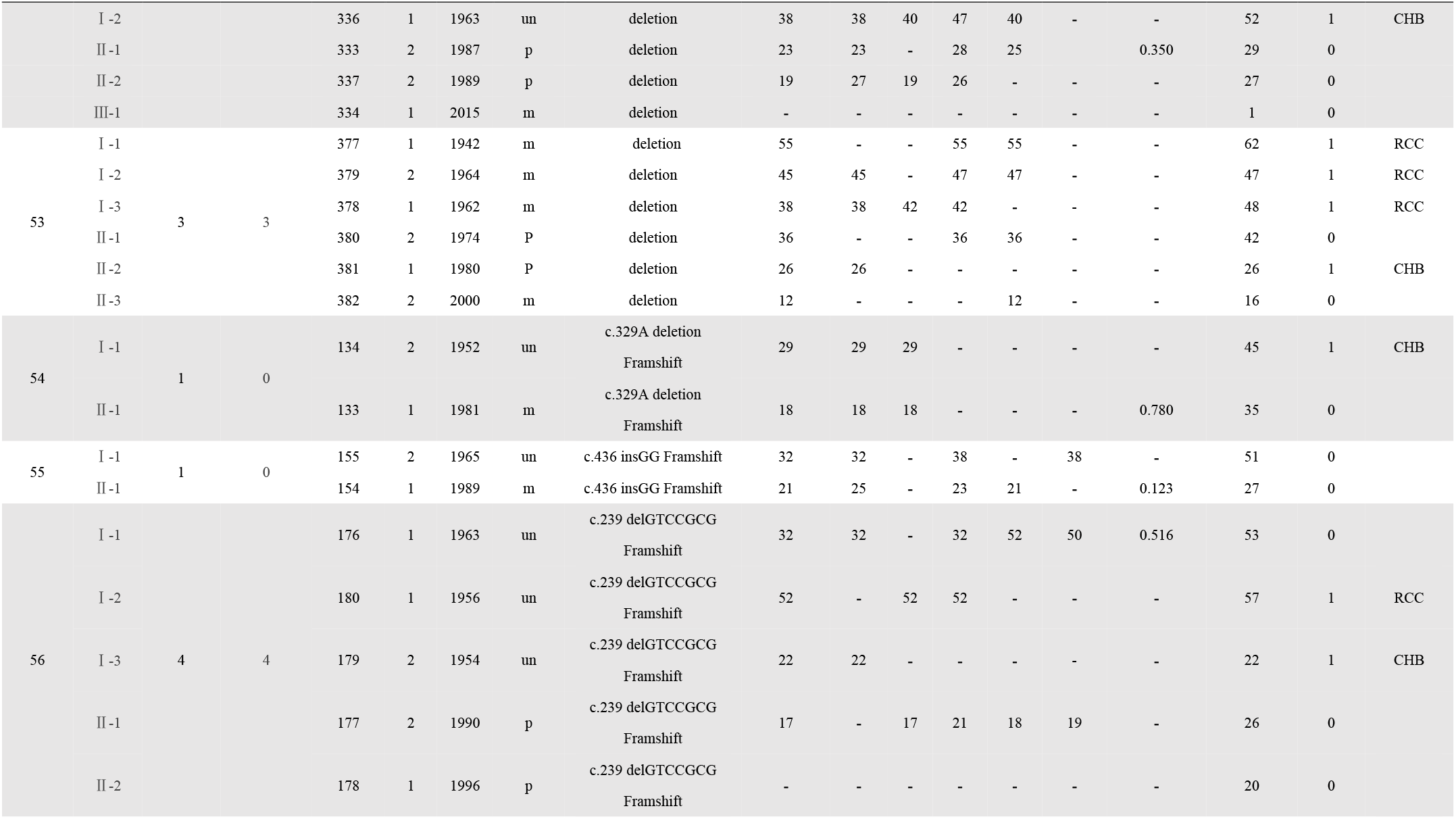
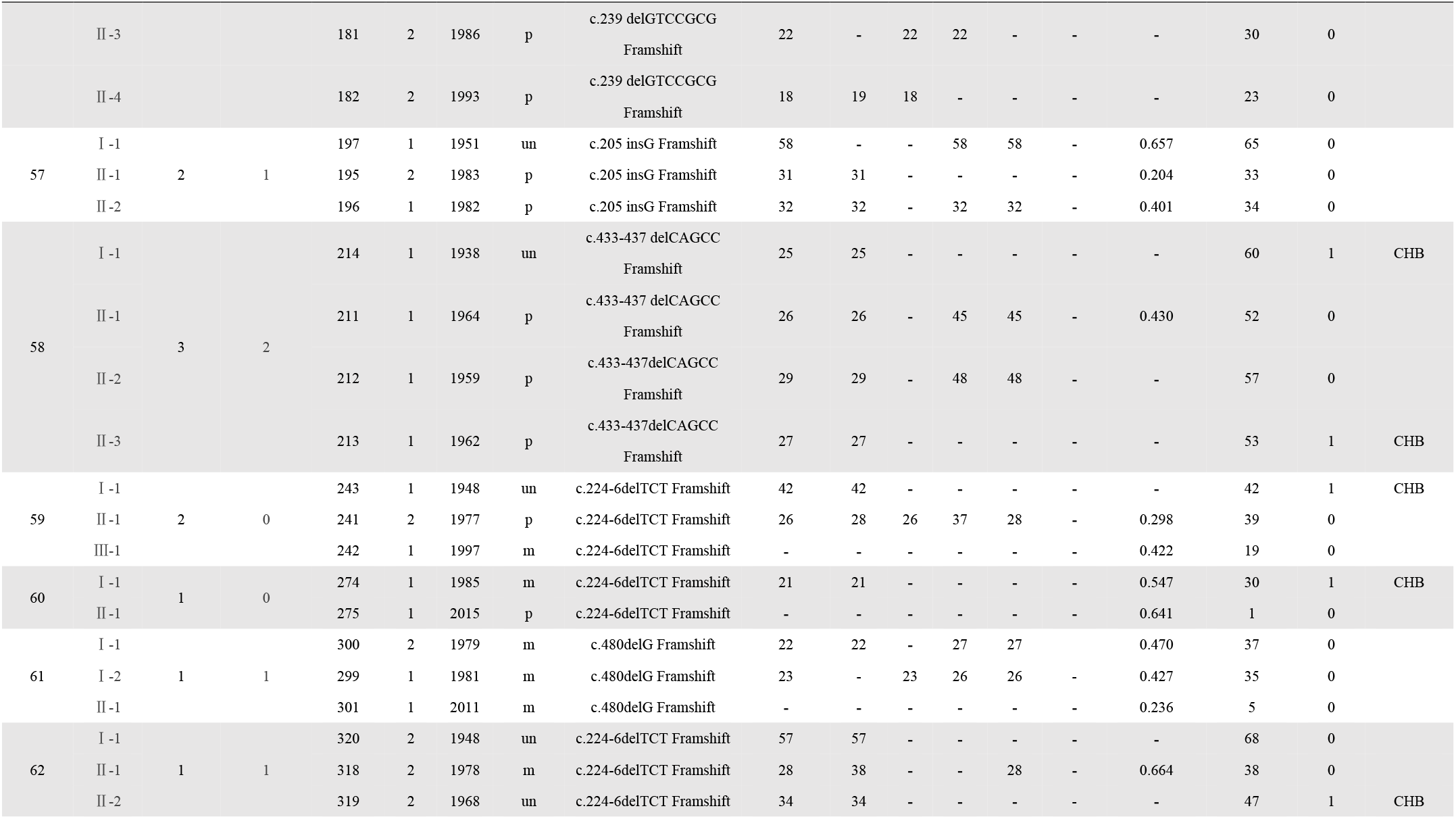
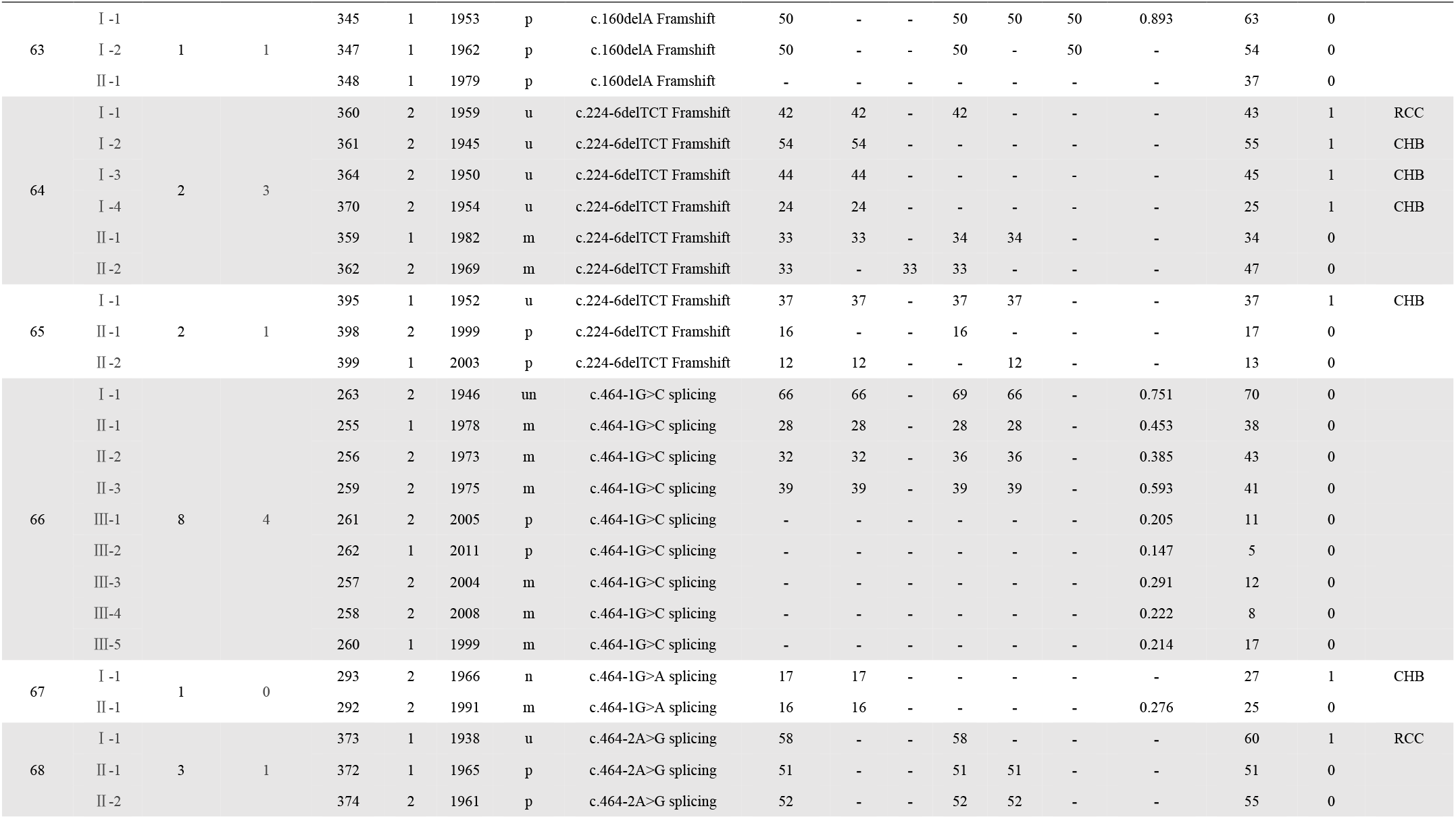
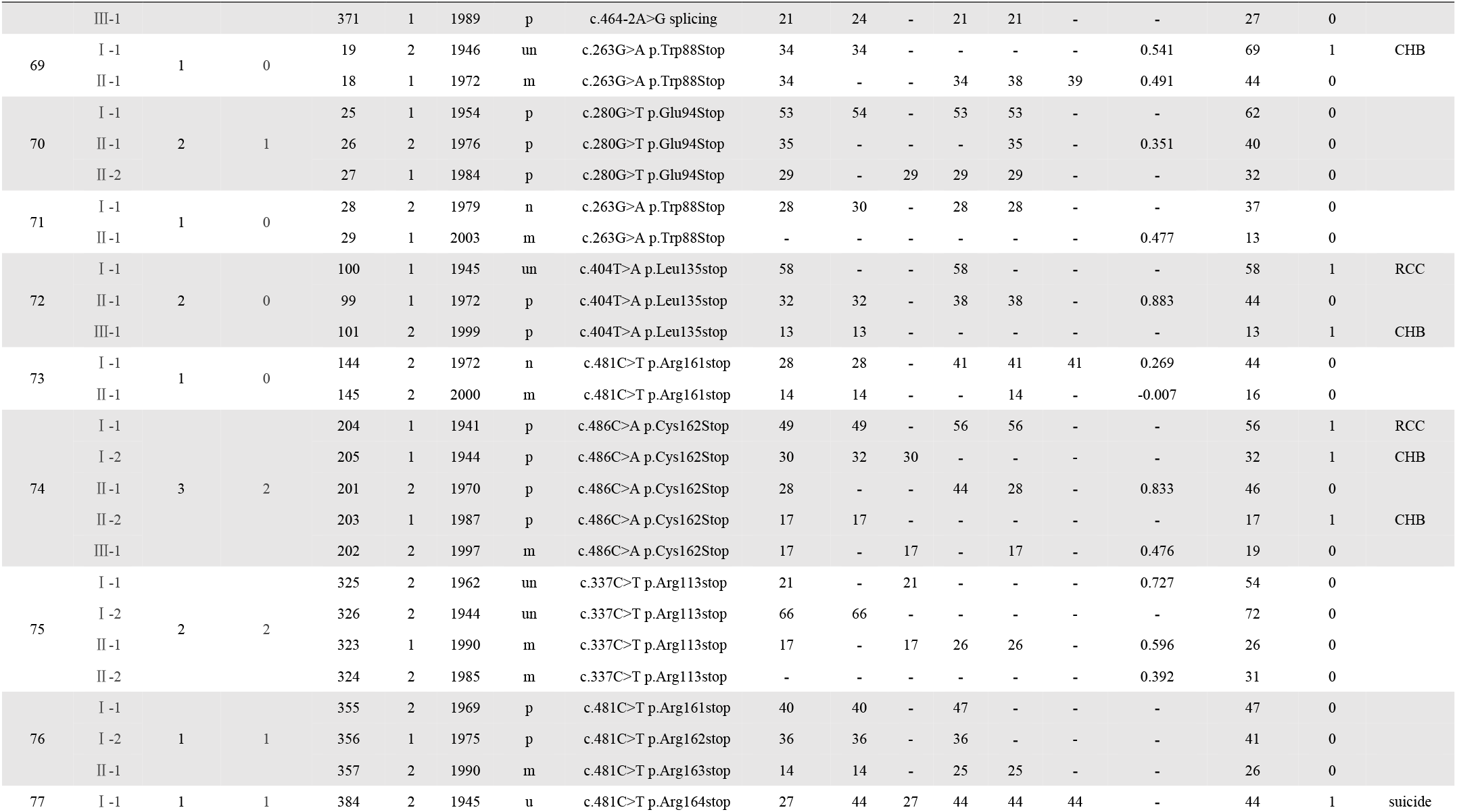
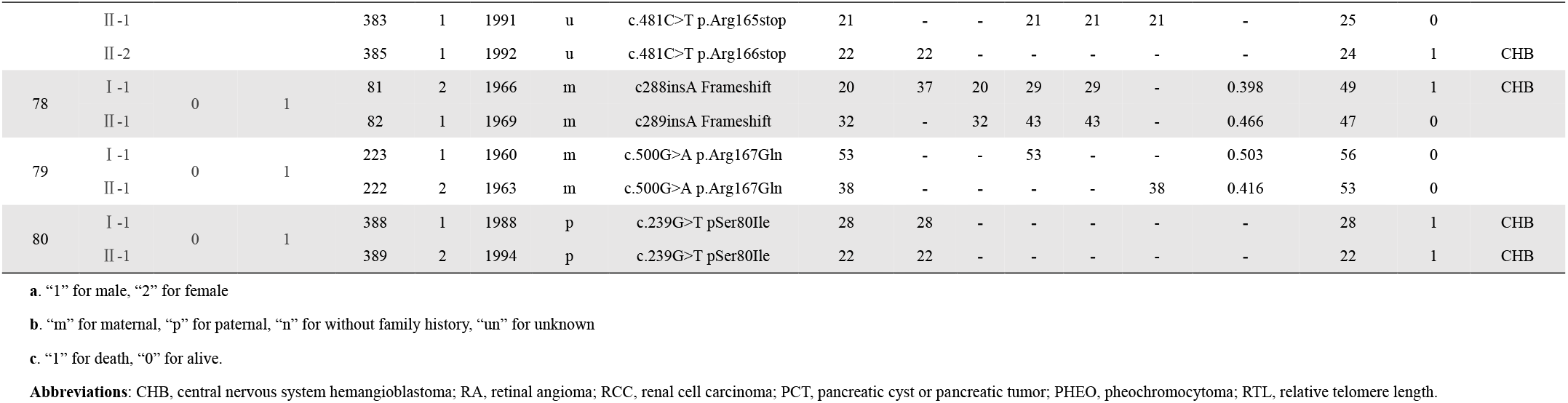
Clinical and genetic characteristics of VHL patients involved in this study

**Table S2.** Genetics anticipation in affected and non-affected VHL patients with CPH model

| Parameter | Estimated (Robust SE) | Wald p value |
| --- | --- | --- |
| Generation | 1.184 (0.142) | <0.001 |
| Gender | 0.116 (0.139) | 0.400 |
| Mutation type | 0.188 (0.139) | 0.180 |

Abbreviations: CPH, Cox proportional hazards.

**Table S3.** Genetic anticipation in different organs in affected parents-children pairs

| **Organ** | **Pairs** | **Mean onset age (y)** |  | **MOAD(y)** | **P**<br>**(paired t test)** |
| --- | --- | --- | --- | --- | --- |
|  |  | **Parent** | **Children** |  |  |
| **CHB** | 62 | 42.3±13.3 | 25.2±9.4 | 17.2 | <0.001 |
| **RCC** | 29 | 52.5±9.6 | 31.4±8.8 | 21.1 | <0.001 |
| **PCT** | 25 | 54.0±10.5 | 28.3±9.1 | 25.8 | <0.001 |
| **RA** | 10 | 35.8±12.4 | 16.5±3.1 | 19.3 | <0.001 |
| **PHEO** | 5 | 52.8±5.8 | 18.2±6.6 | 34.6 | 0.0012 |

Abbreviations: MOAD, mean of onset age difference; CHB, central nervous system hemangioblastoma; RA, retinal angioma; RCC, renal cell carcinoma; PCT, pancreatic cyst or pancreatic tumor; PHEO, pheochromocytoma.

**Table S4.** Telomere length between sibling pairs

|  | Onset age (O>Y) | Onset age (O≤Y) | Overall |
| --- | --- | --- | --- |
| **Age-adjusted RTL**<br>**(O>Y)** | 8 | 5 | 13 |
| **Age-adjusted RTL**<br>**(O≤Y)** | 3 | 3 | 6 |
| **Overall** | 11 | 8 | 19 |

Abbreviations: O, Older; Y, Younger; RTL, relative telomere length.

